# Metagenomic analysis reveals the signature of gut microbiota associated with human chronotypes

**DOI:** 10.1101/2021.03.16.435653

**Authors:** Shaqed Carasso, Bettina Fishman, Liel Stelmach Lask, Tamar Shochat, Naama Geva-Zatorsky, Eran Tauber

**Affiliations:** Department of Cell Biology and Cancer Science, Rappaport Faculty of Medicine Rappaport Technion Integrated Cancer Center (TICC), Haifa 3525422, Israel; Department of Evolutionary and Environmental Biology and Institute of Evolution, University of Haifa, Haifa 3498838, Israel; Cheryl Spencer Department of Nursing, Faculty of Social Welfare and Health Sciences, University of Haifa, Haifa 3498838, Israel; Canadian Institute for Advanced Research (CIFAR), MaRS Centre, West Tower 661 University Ave., Suite 505 Toronto, ON M5G 1M1, Canada

**Keywords:** Chronotype, circadian clock, gut microbiome, metagenomics, sleep

## Abstract

Patterns of diurnal activity differ substantially between individuals, with early risers and late sleepers being examples of opposite chronotypes. Growing evidence suggests that the late chronotype significantly impacts the risk of developing mood disorders, obesity, diabetes, and other chronic diseases. Despite the vast potential of utilizing chronotype information for precision medicine, those factors that shape chronotypes remain poorly understood. Here, we assessed whether the various chronotypes are associated with different gut microbiome compositions. Using metagenomic sequencing analysis, we established a distinct signature associated with chronotype based on two bacterial genera, *Alistipes* (elevated in “larks”) and *Lachnospira* (elevated in “owls”). We identified three metabolic pathways associated with the early chronotype, and linked distinct dietary patterns with different chronotypes. Our work demonstrates an association between the gut microbiome and chronotype and may represent the first step towards developing dietary interventions aimed at ameliorating the deleterious health correlates of the late chronotype.

## Introduction

Numerous biological, mental, and behavioral functions present circadian oscillations that are orchestrated by a central pacemaker in the brain. While the periodicity of these rhythms under natural conditions is uniform (24 h), their phase shows considerable inter-individual variability. Phase variation is often manifested as the tendency of individuals to be active during specific times of the day, referred to as chronotype. There is a genetic predisposition for chronotype,^1^ which changes over the course of development and determines individual variation in the diurnal timing of various functions, such as sleep, cognitive and physical activities.^2^

Early studies in rodents demonstrated that high-fat diets altered both circadian rhythm locomotor output and the expression of circadian clock genes.^3^ Subsequently, the effect of diet on the circadian clock was found to be mediated by the gut microbiome, which has been shown to regulate metabolic processes and modulate brain functions and behavior via immune, endocrine and neural pathways via the gut-brain axis.^4^

Studies conducted in the last decade revealed diurnal oscillations in gut microbiome function and composition.^5–7^ Using murine models, these studies found that about 20% of the gut microbiota showed diurnal oscillations in terms of composition, as well as in associated functions and metabolites. These microbiota oscillations are linked to the circadian clock, as demonstrated in clock mutant *Per1/2*^-/-5^ or *Bmal1* knockout mice,^8^ in which these microbiota rhythms were abolished. Furthermore, disruption of circadian rhythms by phase reversal of the light-dark cycle (combined with a high-fat, high-sugar diet) led to intestinal dysbiosis.^9^ Similarly, perturbations of the circadian rhythm in the host impacted the intestinal microbiota. ^5, 10^ On the other hand, ablation of the microbiome by antibiotic treatment disrupted normal transcriptional circadian oscillation of the host, apparently via reprogramming of the chromatin state.^11^

Studies on diurnal oscillation of the gut microbiota in humans are clearly more challenging. Pioneering studies with a very limited number of subjects corroborated diurnal oscillation of the microbiota.^5, 12^ A more recent study demonstrated microbiome rhythmicity in two very large cohorts using collection time-stamp data.^13^ As these cohorts included type-2 diabetes subjects, it was possible to identify several rhythmic taxa that were associated with the disease and which could serve as powerful predictors and potential nodes for future interventions.

Although previous findings alluded to bidirectional associations between gut microbiota and the circadian clock, it remains unclear whether variation in chronotype is also linked to specific microbiota compositions. Here, we considered this link using metagenomic sequencing of individuals assigned distinct chronotypes.

## Materials and Methods

This study was approved by the local ethics committees of the University of Haifa (#283/18). Participants provided written informed consent prior to participation.

The study population consisted of 133 individuals (51% males), evaluated for body mass index (BMI) and medication intake. Of these, only medication-free individuals who do not use an alarm clock on free days were included (N = 91). Chronotype was assessed by computing mid-sleep time on free days (MSF).^2^ Participants’ stool samples were collected using DNA/RNA Shield Fecal Collection tubes (Zymo research). Sample DNA extraction was performed using the PureLink Microbiome DNA Purification Kit (Invitrogen) according to the manufacturer’s instructions. Genomic DNA was sheared to an average size of 300 bp with an M220 ultrasonicator (Covaris, Woburn, MA). Sheared DNA samples were used for paired-end indexed library construction using an Ovation Ultralow library systems V2 kit (NuGEN, San Carlos, CA), according to the manufacturer instructions.

Next-generation sequencing (Paired end, 2×150 bp) was performed by the Genome Research Core, University of Illinois at Chicago, using the Illumina NextSeq500 sequencer. Metagenomic sequencing data was processed with bioBakery workflows utilizing bioBakery 3 tools.^14^ Briefly, sequence data quality control, including removal of human reads, was performed using KneadData. Taxonomic profiles were generated using MetaPhlAn v3.0 and functional profiles were generated with HUMAnN v3.0 using MetaCyc pathway definitions.^15^ Relative proportion data were transformed by centered-log ratio. Microbial communities were compared among the different chronotypes. Primary analysis was performed using MicrobiomeAnalyst^16^ followed by comprehensive analysis using the R packages Phyloseq^17^, Vegan, and MaAsLin2 v1.06. Alpha diversity indices (Chao1 and Shannon) were compared using the Wilcoxon rank sum test. Beta diversity distance matrices (Aitchison) were visualized using principal coordinate analysis (PCA) and compared using the Vegan package’s function ADONIS, a multivariate analysis of variance based on dissimilarity tests. Associations between participant characteristics and their microbial taxa or functional modules were assessed using differential abundance analyses with MaAsLin2. Results were visualized using the ggplot2 R package.

All raw metagenomic data were deposited in the National Center for Biotechnology Information sequence read archive as BioProject accession PRJNA714678.

## Results and Discussion

To investigate changes in the gut microbiome composition of different chronotypes, we collected fecal samples from 91 individuals from across Israel (50 females, mean age ± SD: 33.85 ± 10.95 years; body mass index (BMI): 23.6 ± 3.7).

The distribution of chronotypes is shown in Figure 1A. MSF distribution centered on 4:12 (± 00:15 h (SD)) and did not differ significantly between males and females (Watson-Williams F test, *p* =0.2). For analysis of microbiome composition, participants were divided into three groups based on their MSF as “Early” (n = 24), “Intermediate” (n = 27) and “Late” (n = 40) chronotypes.

**Figure 1.**
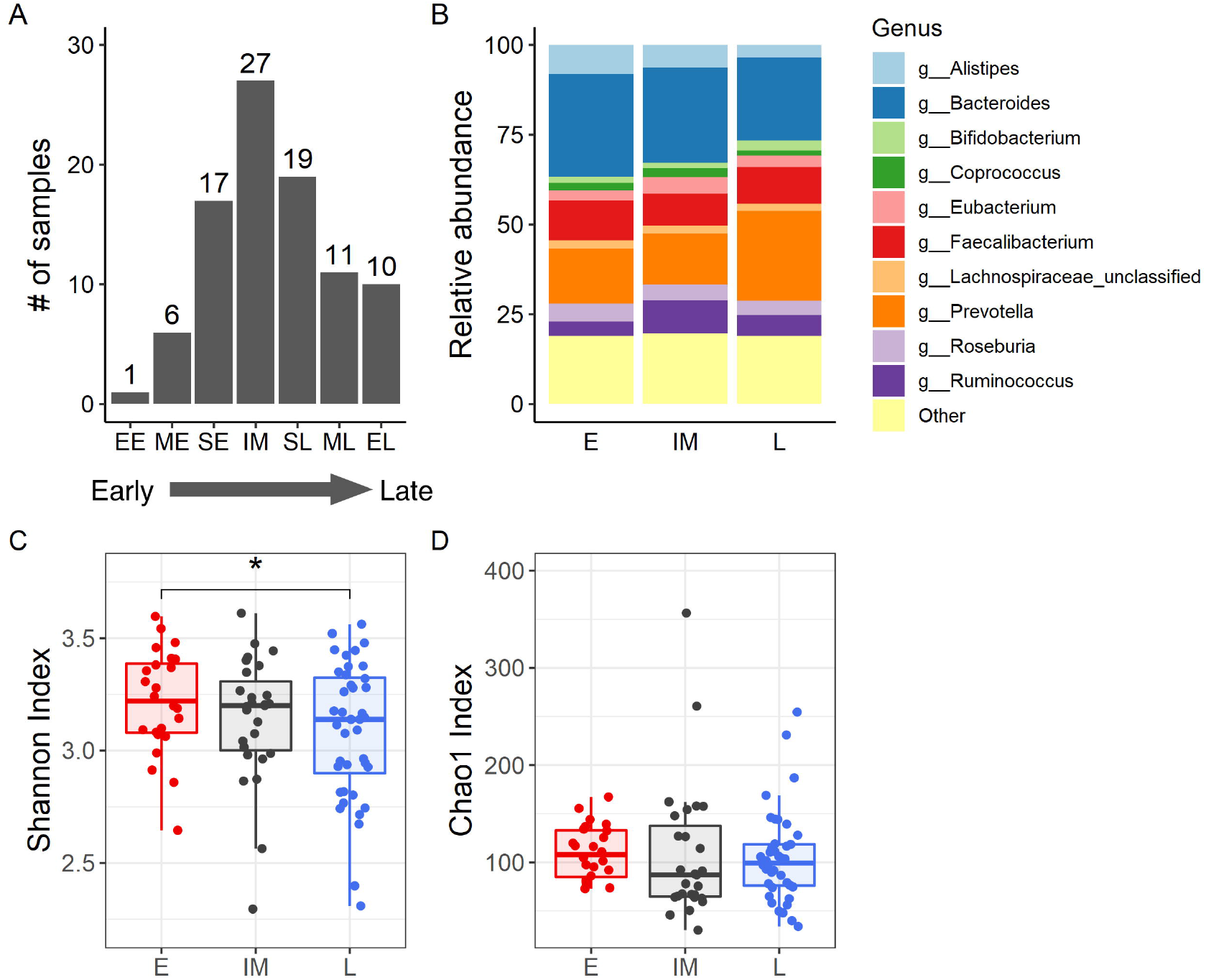
**A.** Chronotype distribution. Chronotypes were divided into 7 categories as Extremely early (EE): <01:29; Moderately early (ME): 1:30-2:30; Slightly early (SE): 2:30-3:30; Intermediate (IM): 3:30-4:30; Slightly late (SL): 4:30-5:30; Moderately late (ML): 5:30-6:30; and Extremely late (EL): > 6:30. **B.** Microbiome composition (genus level) in the different chronotypes. The relative abundance of main taxa in early (E, comprising the pooled EE, ME and SE populations) intermediate (IM) and late (L, comprising the pooled SL, ML and EL populations) chronotype is shown. **C.** Alpha diversity was measured via the Shannon index and **D.** using the Chao1 index, in the early (E) intermediate (IM) and late (L) chronotypes. Data represent the median (line in box), IQR (box), and minimum/maximum (whiskers). Statistical comparisons are shown for separate matched-pairs tests determined with the Kruskal-Wallis rank sum test. *: p<0.05; **: p< 0.01.

DNA from fecal samples was extracted for shotgun metagenomics sequencing. The mean library size was 3,091,458 reads (ranging from 174,934 to 19,829,176 reads). Samples were filtered to include those taxa with a minimum relative abundance of >0.01% and identified in 20% of the samples. The identified taxa in the filtered dataset were distributed into 7 phyla, 14 classes, 18 orders, 30 families, 57 genera, and 123 species. The most ubiquitous genera were the *Bacteroides, Faecalibacterium, Parabacteroides,* and *Eubacterium,* all detected in more than 96% of the samples. The ten most abundant genera are presented in Figure 1B.

We observed a higher α-diversity (Shannon index) in participants from the early than from the late chronotype (p < 0.05, Kruskal-Wallis rank sum test), indicating a less complex microbiota community in the latter. There were no significant differences between early and intermediate or between intermediate and late chronotypes (Figure 1C). Chao1 indices, measuring the richness of the samples, were not significantly different between chronotypes (Figure 1D). It should be acknowledged that Chao1 is dependent on singletons and doubletons that might rise because of sequencing errors. Beta diversity based PCA did not reveal a clear difference between chronotypes (Adonis test, p = 0.375; Supp. Figure 1).

Group differences in relative gut bacteria abundance were tested using a feature-wise association approach, controlling for age, BMI, and gender. Two genera were significantly different in the early and late chronotypes, namely, *Alistipes* (pFDR = 0.2; Figure 2A) and *Lachnospira* (pFDR = 0.2; Figure 2B). Within these genera, two species were significantly different between groups. *Alistipes putredinis* was more abundant in the early chronotype, as compared to both the intermediate and late chronotypes (pFDR = 0.2; Figure 2C). *Lachnospira pectinoschiza* was more abundant in the late than in the early chronotype (pFDR = 0.2; Figure 2D).

**Figure 2.**
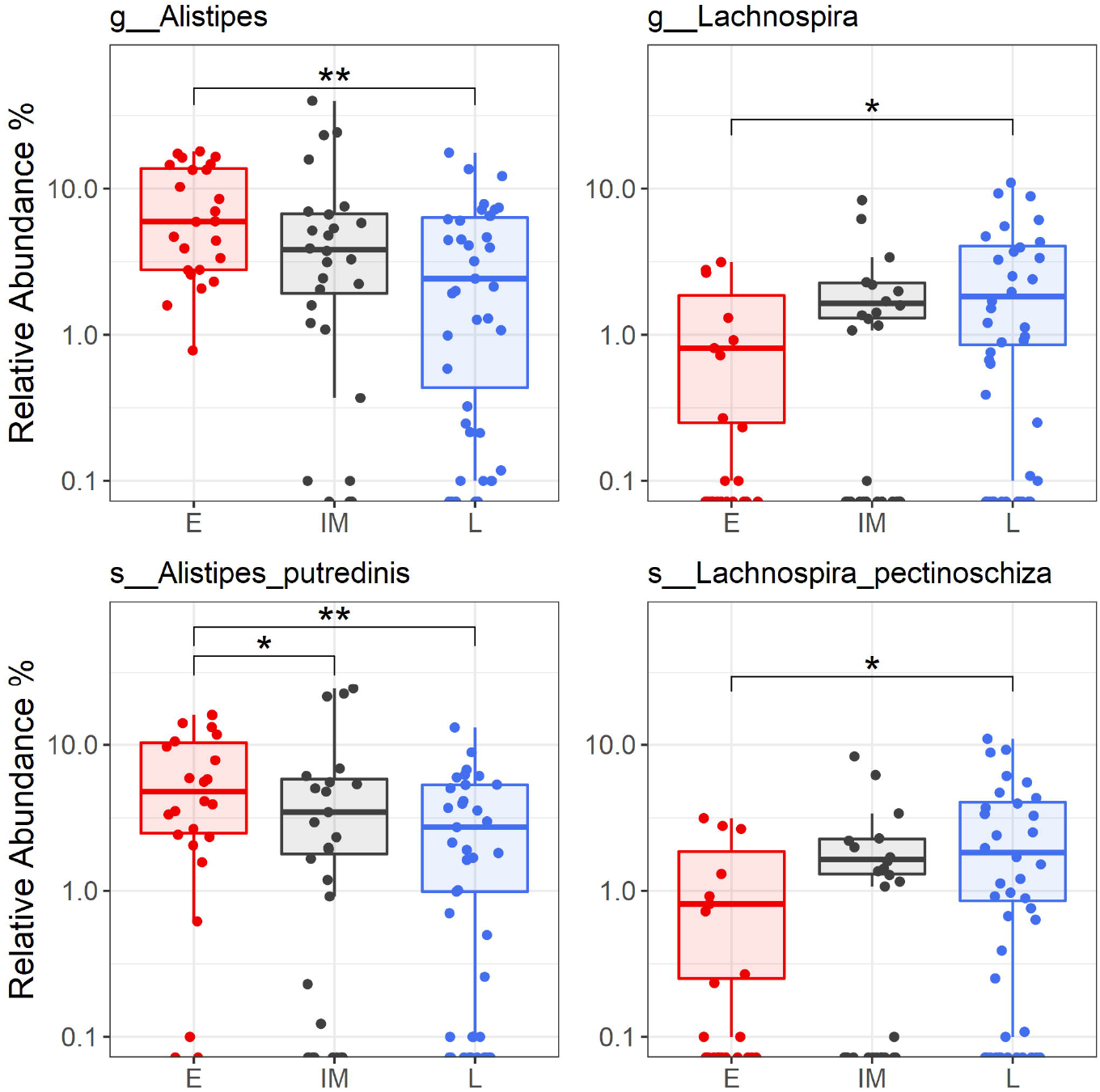
Relative abundances of significantly different taxa in the early (E), intermediate (IM) and late (L) chronotypes. Data represent the median (line in box), IQR (box), and minimum/maximum (whiskers). Statistical comparisons are shown for separate matched-pairs tests determined with the Kruskal-Wallis rank sum test. *: p<0.05; **: p< 0.01.

Reads were also assigned to microbial metabolic pathways and differences in relative abundance were tested. In total, 14 pathways differed significantly among early, intermediate, and late chronotypes (pFDR < 0.2), 11 of which showed differences between intermediate and either early or late chronotypes. The three pathways that differed between early and late chronotypes (Figure 3) included the histidine, purine, and pyrimidine biosynthesis super-pathway (pathway ID PRPP-PWY), of the pyrimidine deoxyribonucleotide *de novo* biosynthesis super-pathway (PWY-7211), and the gluconeogonesis pathway (PWY66-399). All four pathways were more abundant in early, as compared to late chronotypes (Figure 3).

**Figure 3.**
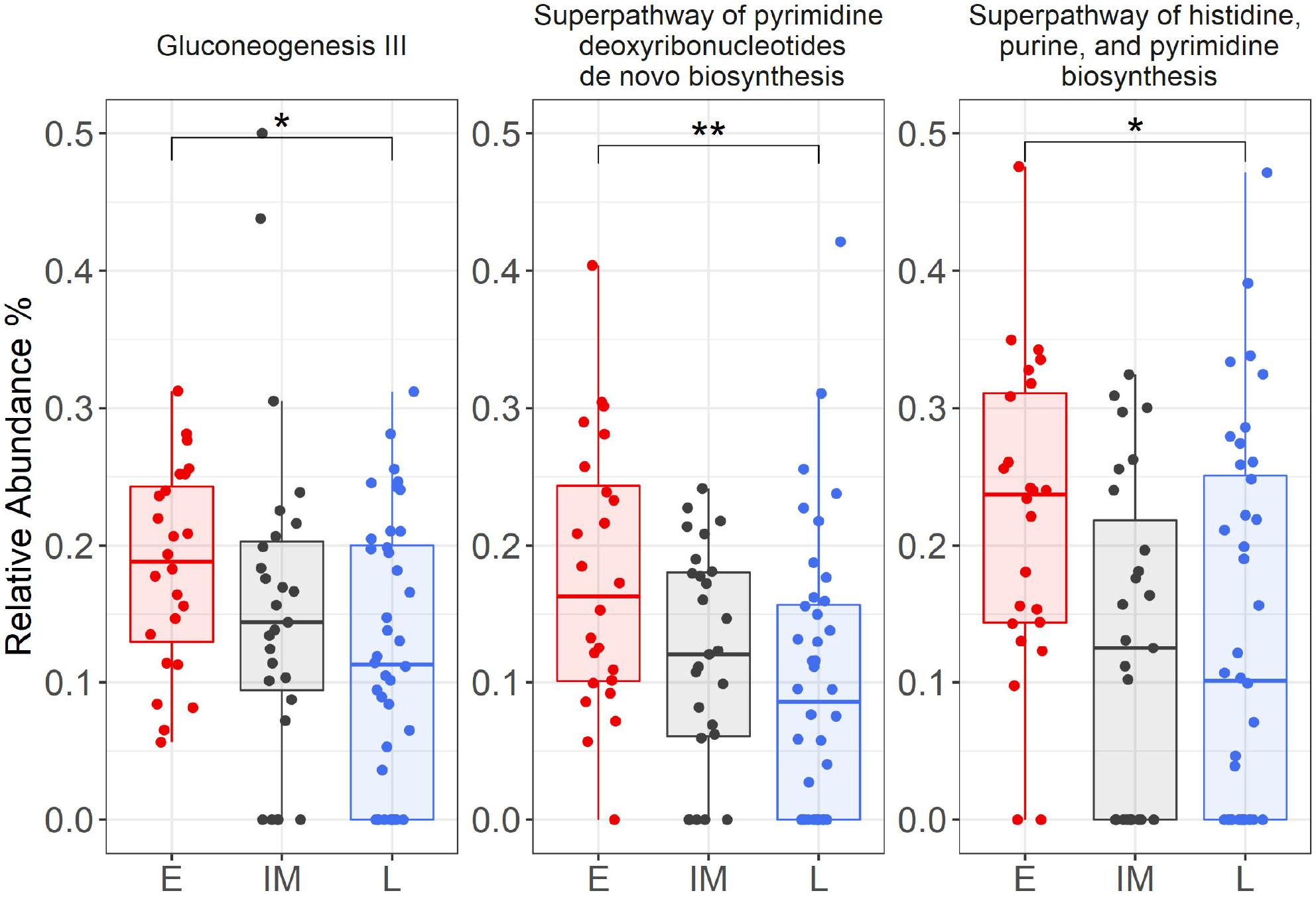
Relative abundances of significantly different metabolic pathways in the early (E), intermediate (IM) and late (L) chronotypes. Data represent the median (line in box), IQR (box), and minimum/maximum (whiskers). Statistical comparisons are shown for separate matched-pairs tests determined with the Kruskal-Wallis rank sum test. *: p<0.05; **: p< 0.01.

In a recent study, the time of defecation was identified as a significant factor contributing to variability in microbiome structure. We, therefore, analyzed this feature in our data. The time of defecation was reported by 48% of participants (N = 44). As expected, the data suggested that early chronotypes were more likely to report defecating in the morning, while late chronotypes reported have defecated in the evening (Supporting information. Figure S2). However, a Chisquare test between time of defecation and chronotype revealed no significant association (X^2^ = 2.56, p =0.635), possibly due to the limited sample size. Importantly, there were no significant associations between time of defecation and relative bacterial abundance (all FDRs were over 0.95).

Since a substantial factor shaping gut microbiome composition is diet,^18^ we analyzed participants’ food consumption. Participants reported weekly consumption frequency of 13 types of food (e.g., beef, chicken, pastry, fruits, and vegetables) and drink (water, sugary or diet drinks). The questionnaire is based on the Israeli National Health and Nutrition (MABAT Cohort) survey (see Supplemental Document 1). Food item counts were analyzed using principal component analysis (PCA) (Figure 4A). The first principal component (PC1, 17.7%) revealed a major dietary difference between participants who consumed a healthy diet (including fruits, vegetables, water), and those with an unhealthy diet (including high-sugar, high fat, sugary drinks). A small but highly significant correlation was found between chronotype and PC1 scores (p < 0.01; Figure 4B). As reported previously,^19^ early chronotypes exhibited higher adherence towards a healthy diet.

**Figure 4.**
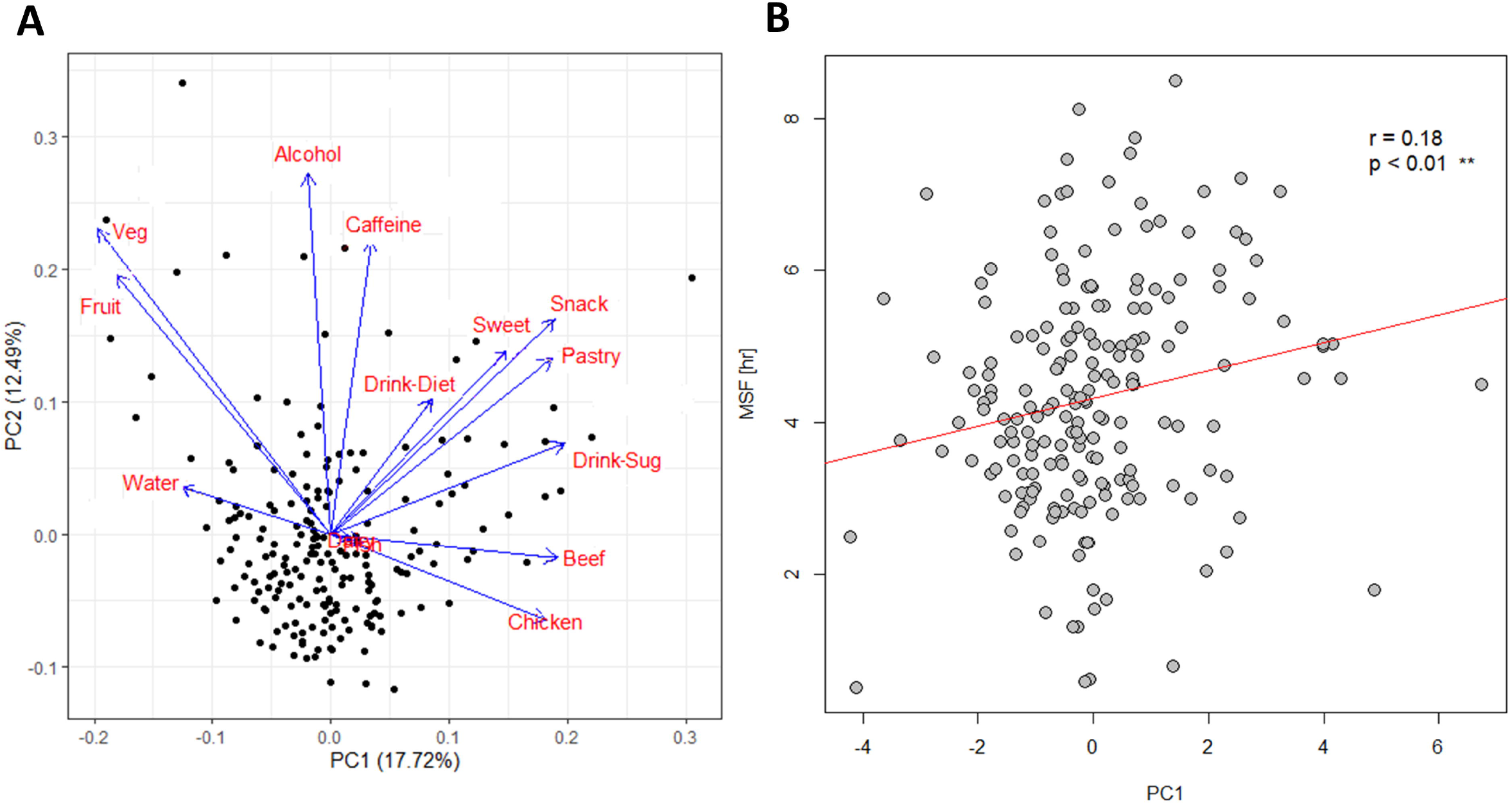
**A.** PCA of food and drink items consumed by the participants. **B.** Correlation between chronotype (MSF) with PC1 scores from the PCA showing a significant trend (p < 0.01). Fruits, vegetables (and water as a drink) are prominent in early chronotypes, while meat, pastry and sugary drinks are associated with late chronotypes.

We next assessed whether the identified differences in bacteria were correlated with participant dietary habits. As shown in Figure 4B, higher consumption of fruits and vegetables was correlated with the early chronotype. Hence, we examined association between vegetable and fruit consumption and the microbiome. The reported number of vegetables and fruits consumed weekly was aggregated and participants were divided into those who consumed ≤ or >10 vegetables and/or fruits. The findings showed that the *Ruminococcaceae* family was more abundant in participants who consumed >10 fruits and vegetables in a week. This finding agrees with previous work showing that high fruit intake is positively associated with *Ruminococcaceae* levels in the microbiome.^20^ No additional associations were observed between microbiome composition and food consumption. An increase in the *Negativicutes* bacterial class was associated with higher BMI. Indeed, increased levels of *Negativicutes* were previously reported in obese individuals with type 2 diabetes.^21^

Interestingly, a recent study that analyzed diurnal rhythms of gut microbiota in a large cohort by unsupervised analysis13 revealed three clusters. While chronotypes were not determined in this study, it is quite possible that chronotype contributed to observed difference between the clusters, given the fact that one of clusters was dominated by *Ruminococcus,* and that many physiological traits (e.g. body weight, time of defecation) and lifestyle considerations (e.g. alcohol consumption) are associated with chronotype.

Overall, our analysis revealed a few consistent differences among chronotypes. The finding that lower α-diversity, a marker of dysbiosis, was observed in late chronotypes, is consistent with growing evidence of increased cardio-metabolic morbidity and mortality risk in this group.^22^ Furthermore, previous studies showed that jet lag in both humans and mice is associated with dysbiosis,^5^ and since late chronotype individuals are predisposed to social jet lag (i.e., misalignment of social and endogenous circadian clocks), the decreased microbial diversity we observed was to be expected.

The association between chronotype and dietary intake is well established. In addition to lower consumption of fruits and vegetables^24^ and less healthy diet intake by late chronotype individuals,^19^ there is a marked difference in meal timing between the chronotypes.^25^ Meal timing was shown to be an important factor impacting microbiota diurnal rhythms,^26^ such that not only dietary composition but also meal timing shapes our microbiota profile. Collado *et al.* analyzed the salivary microbiota, which unlike the gut microbiome, allows for repeated serial sampling over 24 h.^26^ These authors found that delaying the main meal inverted the diurnal rhythm of salivary microbiota diversity. In our study, the timing of meals was not recorded, yet given the established difference between early and late chronotypes in meal timing, it is possible that this difference augmented differences in microbiota composition between these chronotypes.

We identified two bacterial genera that differed between early and late chronotypes. Interestingly, *Alistipes,* which was enriched in early chronotypes, was previously reported as being over-represented in older mice^27^ and humans.^28^ This may explain the well-known observation that chronotype becomes earlier with age, with MSF advancing by nearly two hours between the ages of 20 to 70 years.^29^ Furthermore, *Lachnospira* was more abundant in late chronotypes. In a recent study on human microbiota and eating behavior during the day, a higher abundance of *Lachnospira* was found when a greater percentage of energy was consumed after 2 p.m.^12^. This finding aligns with our data portraying *Lachnospira* as a biomarker of the late chronotype, as energy consumption is expected to be delayed in these individuals.^30^

A limitation of our study is the rather small sample sizes of chronotype sub-group considered, particularly the number of early (E) chronotype participants. Studies with larger cohorts will improve statistical power and may lead to the identification of additional differences between the early and late chronotypes.

Finally, our study identified several metabolic pathways that differed between chronotypes, particularly gluconeogenesis (Figure 3), which is known to be under tight circadian regulation in host cells.^31^ Interestingly, intestinal gluconeogenesis is activated by short-chain fatty acids (SCFAs), particularly butyrate, produced by the fermentation of soluble fibers by gut bacteria.^32^ Emerging evidence suggests that SCFAs serve as signaling molecules between the gut microbiota and host metabolism.^33^ Furthermore, the *Lachnospira* species identified here as a marker of the late chronotype is a member of a family known to synthesize butyrate.^25^ In mice, robust diurnal oscillation was observed for both *Lachnospira* species as well as butyrate levels.^6^ Importantly, SCFAs were recently implicated in sleep regulation in rodents^34^ and humans,^35, 36^ indicating a possible link between gluconeogenesis, the microbiota and host chronotype, which is mediated by SCFAs. This putative link merits future exploration.

## ACKNOWLEDGMENTS

We thank two anonymous reviewers for their insightful comments. We thank Bengisu Sezen Subaşı for her technical assistance, and the N.G-Z. lab for discussions and help. This study was supported by a research grant from the Technion-University of Haifa joint projects funded by the Milgrom family to E.T., T.S and N. G-Z. and the Israel Science Foundation (grant 1737/17 to E.T.). N.G-Z. was supported by CIFAR (Azrieli-Global Scholar), the Technion, “Hanasi”, “Cathedra”, “Alon” and “Horev” fellowships and the Human Frontiers Science Program (HFSP CDA00025/2019-C).

## AUTHORS’ CONTRIBUTIONS

E.T., T.S. and N.G-Z. conceived the project, N.G-Z. contributed to the design and implementation of the research, B.F. and T.S. recruited study participants, B.F. carried out data collection (stool samples and questionnaires) and ran the experiments, and S.C. carried out the bioinformatics analysis. All authors discussed the results and contributed to the final manuscript.

## Non-standard abbreviations

MSF: mid-sleep time on free days
FDR: false discovery rate
PCA: principal component analysis

## References

1. Jones, S. E., Lane, J. M., Wood, A. R., van Hees, V. T., Tyrrell, J., Beaumont, R. N., Jeffries, A. R., Dashti, H. S., Hillsdon, M., Ruth, K. S., Tuke, M. A., Yaghootkar, H., Sharp, S. A., Jie, Y., Thompson, W. D., Harrison, J. W., Dawes, A., Byrne, E. M., Tiemeier, H., Allebrandt, K. V., Bowden, J., Ray, D. W., Freathy, R. M., Murray, A., Mazzotti, D. R., Gehrman, P. R., Lawlor, D. A., Frayling, T. M., Rutter, M. K., Hinds, D. A., Saxena, R., and Weedon, M. N. Genome-wide association analyses of chronotype in 697,828 individuals provides insights into circadian rhythms. Nat. Commun. 2019; 10:343

2. Roenneberg, T., Wirz-Justice, A., and Merrow, M. Life between clocks: Daily temporal patterns of human chronotypes. J. Biol. Rhythms 2003; 18:80–90

3. Kohsaka, A., Laposky, A. D., Ramsey, K. M., Estrada, C., Joshu, C., Kobayashi, Y., Turek, F. W., and Bass, J. High-Fat Diet Disrupts Behavioral and Molecular Circadian Rhythms in Mice. Cell Metab. 2007; 6:414–421

4. Mu, C., Yang, Y., and Zhu, W. Gut microbiota: The brain peacekeeper. Front. Microbiol. 2016;7

5. Thaiss, C. A., Zeevi, D., Levy, M., Zilberman-Schapira, G., Suez, J., Tengeler, A. C., Abramson, L., Katz, M. N., Korem, T., Zmora, N., Kuperman, Y., Biton, I., Gilad, S., Harmelin, A., Shapiro, H., Halpern, Z., Segal, E., and Elinav, E. Transkingdom control of microbiota diurnal oscillations promotes metabolic homeostasis. Cell 2014; 159:514–529

6. Leone, V., Gibbons, S. M., Martinez, K., Hutchison, A. L., Huang, E. Y., Cham, C. M., Pierre, J. F., Heneghan, A. F., Nadimpalli, A., Hubert, N., Zale, E., Wang, Y., Huang, Y., Theriault, B., Dinner, A. R., Musch, M. W., Kudsk, K. A., Prendergast, B. J., Gilbert, J. A., and Chang, E. B. Effects of diurnal variation of gut microbes and high-fat feeding on host circadian clock function and metabolism. Cell Host Microbe 2015; 17:681–689

7. Zarrinpar, A., Chaix, A., Yooseph, S., and Panda, S. Diet and Feeding Pattern Affect the Diurnal Dynamics of the Gut Microbiome. Cell Metab. 2014; 20:1006–1017

8. Liang, X., Bushman, F. D., and FitzGerald, G. A. Rhythmicity of the intestinal microbiota is regulated by gender and the host circadian clock. Proc. Natl. Acad. Sci. 2015; 112:10479–10484

9. Voigt, R. M., Forsyth, C. B., Green, S. J., Mutlu, E., Engen, P., Vitaterna, M. H., Turek, F. W., and Keshavarzian, A. Circadian Disorganization Alters Intestinal Microbiota. PLoS One 2014; 9:e97500

10. Deaver, J. A., Eum, S. Y., and Toborek, M. Circadian Disruption Changes Gut Microbiome Taxa and Functional Gene Composition. Front. Microbiol. 2018; 0:737

11. Thaiss, C. A., Levy, M., Korem, T., Dohnalová, L., Shapiro, H., Jaitin, D. A., David, E., Winter, D. R., Gury-BenAri, M., Tatirovsky, E., Tuganbaev, T., Federici, S., Zmora, N., Zeevi, D., Dori-Bachash, M., Pevsner-Fischer, M., Kartvelishvily, E., Brandis, A., Harmelin, A., Shibolet, O., Halpern, Z., Honda, K., Amit, I., Segal, E., and Elinav, E. Microbiota Diurnal Rhythmicity Programs Host Transcriptome Oscillations. Cell 2016; 167:1495–1510.e12

12. Kaczmarek, J. L., Musaad, S. M., and Holscher, H. D. Time of day and eating behaviors are associated with the composition and function of the human gastrointestinal microbiota. Am. J. Clin. Nutr. 2017; 106:ajcn156380

13. Reitmeier, S., Kiessling, S., Clavel, T., List, M., Almeida, E. L., Ghosh, T. S., Neuhaus, K., Grallert, H., Linseisen, J., Skurk, T., Brandl, B., Breuninger, T. A., Troll, M., Rathmann, W., Linkohr, B., Hauner, H., Laudes, M., Franke, A., Le Roy, C. I., Bell, J. T., Spector, T., Baumbach, J., O’Toole, P. W., Peters, A., and Haller, D. Arrhythmic Gut Microbiome Signatures Predict Risk of Type 2 Diabetes. Cell Host Microbe 2020; 28:258–272.e6

14. McIver, L. J., Abu-Ali, G., Franzosa, E. A., Schwager, R., Morgan, X. C., Waldron, L., Segata, N., and Huttenhower, C. BioBakery: A meta’omic analysis environment. Bioinformatics 2018; 34:1235–1237

15. Caspi, R., Billington, R., Fulcher, C. A., Keseler, I. M., Kothari, A., Krummenacker, M., Latendresse, M., Midford, P. E., Ong, Q., Ong, W. K., Paley, S., Subhraveti, P., and Karp, P. D. The MetaCyc database of metabolic pathways and enzymes. Nucleic Acids Res. 2018; 46:D633–D639

16. Dhariwal, A., Chong, J., Habib, S., King, I. L., Agellon, L. B., and Xia, J. MicrobiomeAnalyst: A web-based tool for comprehensive statistical, visual and metaanalysis of microbiome data. Nucleic Acids Res. 2017; 45:W180–W188

17. McMurdie, P. J. and Holmes, S. Shiny-phyloseq: Web application for interactive microbiome analysis with provenance tracking. Bioinformatics 2015; 31:282–283

18. Rothschild, D., Weissbrod, O., Barkan, E., Kurilshikov, A., Korem, T., Zeevi, D., Costea, P. I., Godneva, A., Kalka, I. N., Bar, N., Shilo, S., Lador, D., Vila, A. V., Zmora, N., Pevsner-Fischer, M., Israeli, D., Kosower, N., Malka, G., Wolf, B. C., Avnit-Sagi, T., Lotan-Pompan, M., Weinberger, A., Halpern, Z., Carmi, S., Fu, J., Wijmenga, C., Zhernakova, A., Elinav, E., and Segal, E. Environment dominates over host genetics in shaping human gut microbiota. Nature 2018; 555:210–215

19. Maukonen, M., Kanerva, N., Partonen, T., Kronholm, E., Konttinen, H., Wennman, H., and Männistö, S. The associations between chronotype, a healthy diet and obesity. Chronobiol. Int. 2016; 33:972–981

20. Jiang, Z., Sun, T., He, Y., Gou, W., Zuo, L., Fu, Y., Miao, Z., Shuai, M., Xu, F., Xiao, C., Liang, Y., Wang, J., Xu, Y., Jing, L., Ling, W., Zhou, H., Chen, Y., and Zheng, J.-S. Dietary fruit and vegetable intake, gut microbiota, and type 2 diabetes: results from two large human cohort studies. BMC Med. 2020 181 2020; 18:1–11

21. Ahmad, A., Yang, W., Chen, G., Shafiq, M., Javed, S., Zaidi, S. S. A., Shahid, R., Liu, C., and Bokhari, H. Analysis of gut microbiota of obese individuals with type 2 diabetes and healthy individuals. PLoS One 2019; 14

22. Knutson, K. L. and von Schantz, M. Associations between chronotype, morbidity and mortality in the UK Biobank cohort. Chronobiol. Int. 2018; 35:1045–1053

23. Mazri, F. H., Manaf, Z. A., Shahar, S., and Ludin, A. F. M. The Association between Chronotype and Dietary Pattern among Adults: A Scoping Review. Int. J. Environ. Res. Public Health 2020; 17

24. Patterson, F., Malone, S. K., Lozano, A., Grandner, M. A., and Hanlon, A. L. Smoking, Screen-Based Sedentary Behavior, and Diet Associated with Habitual Sleep Duration and Chronotype: Data from the UK Biobank. Ann. Behav. Med. 2016; 50:715–726

25. N, S.-M., S, S., K, M., H, O., Y, T., S, S., K, Y., and K, S. The midpoint of sleep is associated with dietary intake and dietary behavior among young Japanese women. Sleep Med. 2011; 12:289–294

26. Collado, M. C., Engen, P. A., Bandín, C., Cabrera-Rubio, R., Voigt, R. M., Green, S. J., Naqib, A., Keshavarzian, A., Scheer, F. A. J. L., and Garaulet, M. Timing of food intake impacts daily rhythms of human salivary microbiota: a randomized, crossover study. FASEB J. 2018; 32:2060

27. Langille, M. G. I., Meehan, C. J., Koenig, J. E., Dhanani, A. S., Rose, R. A., Howlett, S. E., and Beiko, R. G. Microbial shifts in the aging mouse gut Morgan. Microbiome 2014; 2:50

28. Claesson, M. J., Jeffery, I. B., Conde, S., Power, S. E., O’connor, E. M., Cusack, S., Harris, H. M. B., Coakley, M., Lakshminarayanan, B., O’sullivan, O., Fitzgerald, G. F., Deane, J., O’connor, M., Harnedy, N., O’connor, K., O’mahony, D., Van Sinderen, D., Wallace, M., Brennan, L., Stanton, C., Marchesi, J. R., Fitzgerald, A. P., Shanahan, F., Hill, C., Paul Ross, R., and O’toole, P. W. Gut microbiota composition correlates with diet and health in the elderly. Nature 2012; 488:178–184

29. Fischer, D., Lombardi, D. A., Marucci-Wellman, H., and Roenneberg, T. Chronotypes in the US – Influence of age and sex. PLoS One 2017; 12

30. Xiao, Q., Garaulet, M., and Scheer, F. A. J. L. Meal timing and obesity: interactions with macronutrient intake and chronotype. Int. J. Obes. 2019; 43:1701–1711

31. Panda, S., Antoch, M. P., Miller, B. H., Su, A. I., Schook, A. B., Straume, M., Schultz, P. G., Kay, S. A., Takahashi, J. S., and Hogenesch, J. B. Coordinated transcription of key pathways in the mouse by the circadian clock. Cell 2002; 109:307–320

32. De Vadder, F., Kovatcheva-Datchary, P., Goncalves, D., Vinera, J., Zitoun, C., Duchampt, A., Bäckhed, F., and Mithieux, G. Microbiota-Generated Metabolites Promote Metabolic Benefits via Gut-Brain Neural Circuits. Cell 2014; 156:84–96

33. Morrison, D. J. and Preston, T. Formation of short chain fatty acids by the gut microbiota and their impact on human metabolism. Gut Microbes 2016; 7:189

34. Szentirmai, É., Millican, N. S., Massie, A. R., and Kapás, L. Butyrate, a metabolite of intestinal bacteria, enhances sleep. Sci. Reports 2019 91 2019; 9:1–9

35. Magzal, F., Even, C., Haimov, I., Agmon, M., Asraf, K., Shochat, T., and Tamir, S. Associations between fecal short-chain fatty acids and sleep continuity in older adults with insomnia symptoms. Sci. Rep. 2021; 11

36. Heath, A.-L. M., Haszard, J. J., Galland, B. C., Lawley, B., Rehrer, N. J., Drummond, L. N., Sims, I. M., Taylor, R. W., Otal, A., Taylor, B., and Tannock, G. W. Association between the faecal short-chain fatty acid propionate and infant sleep. Eur. J. Clin. Nutr. 2020 749 2020; 74:1362–1365

