## Supporting Information for "Metagenomic analysis reveals the signature of gut microbiota associated with human chronotypes"

**Supplementary Figure S1.** Beta diversity (Aitchison distance) between groups. PCA plot of taxonomic features of early (red), intermediate (black) and late (blue) chronotype participants.

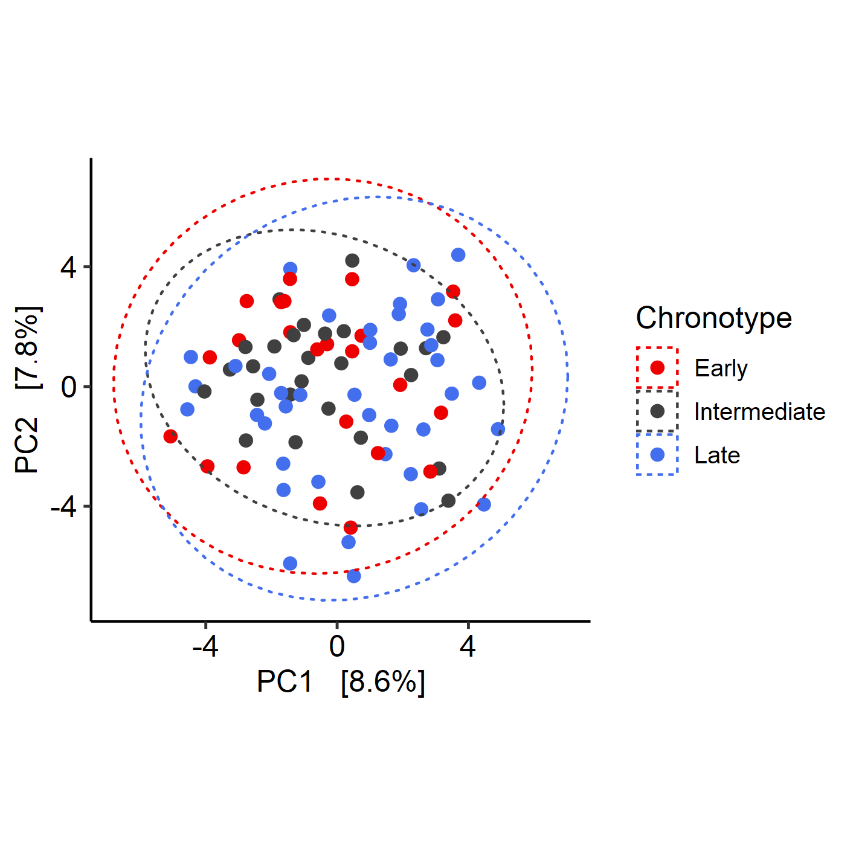

**Supplementary Figure S2.** Distribution of participants’ times of defecation among the early (E), intermediate (IM) and late (L) chronotypes.

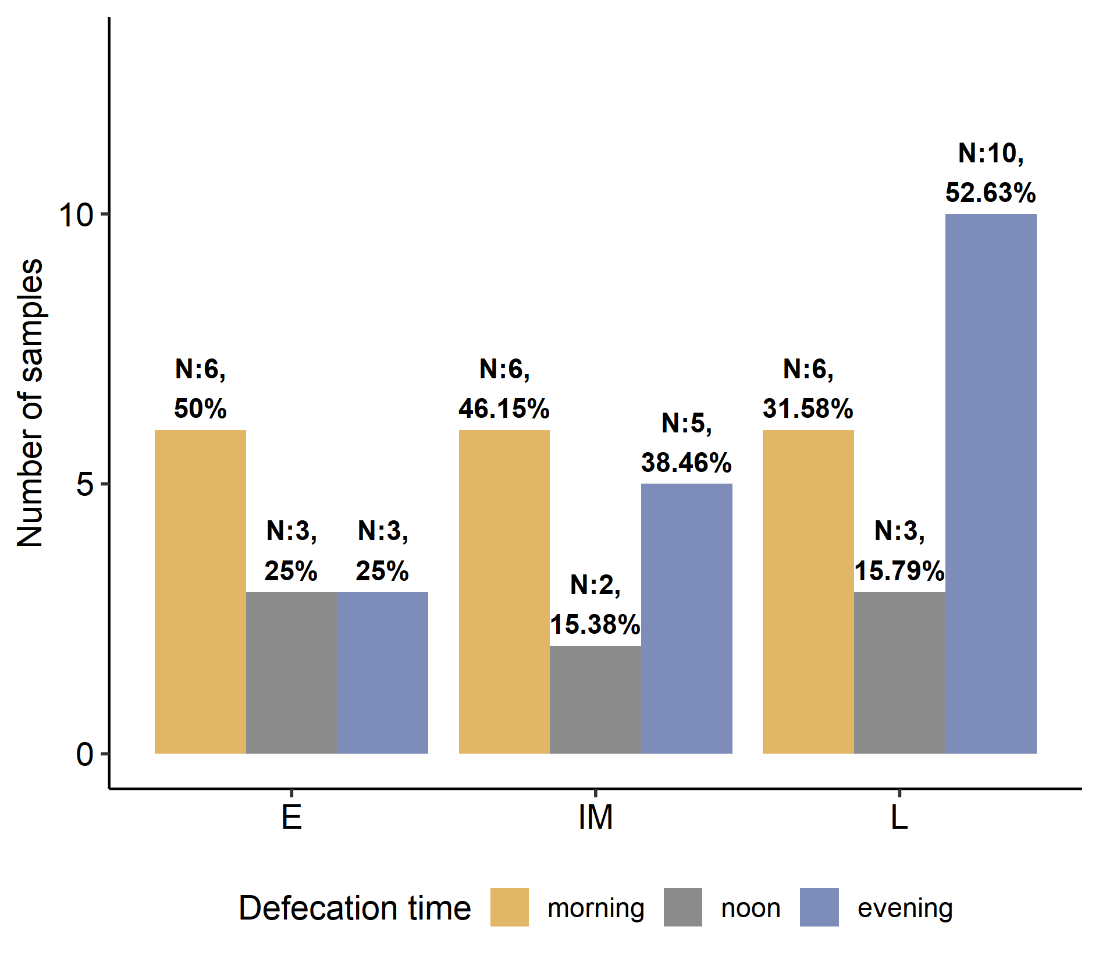

**Supplementary Figure S3.** Relative abundances of the *Negativicutes* bacterial class against BMI, according to chronotype (early (E, red) intermediate (IM, black) and late (L, blue) ). The black line represents the linear regression line.

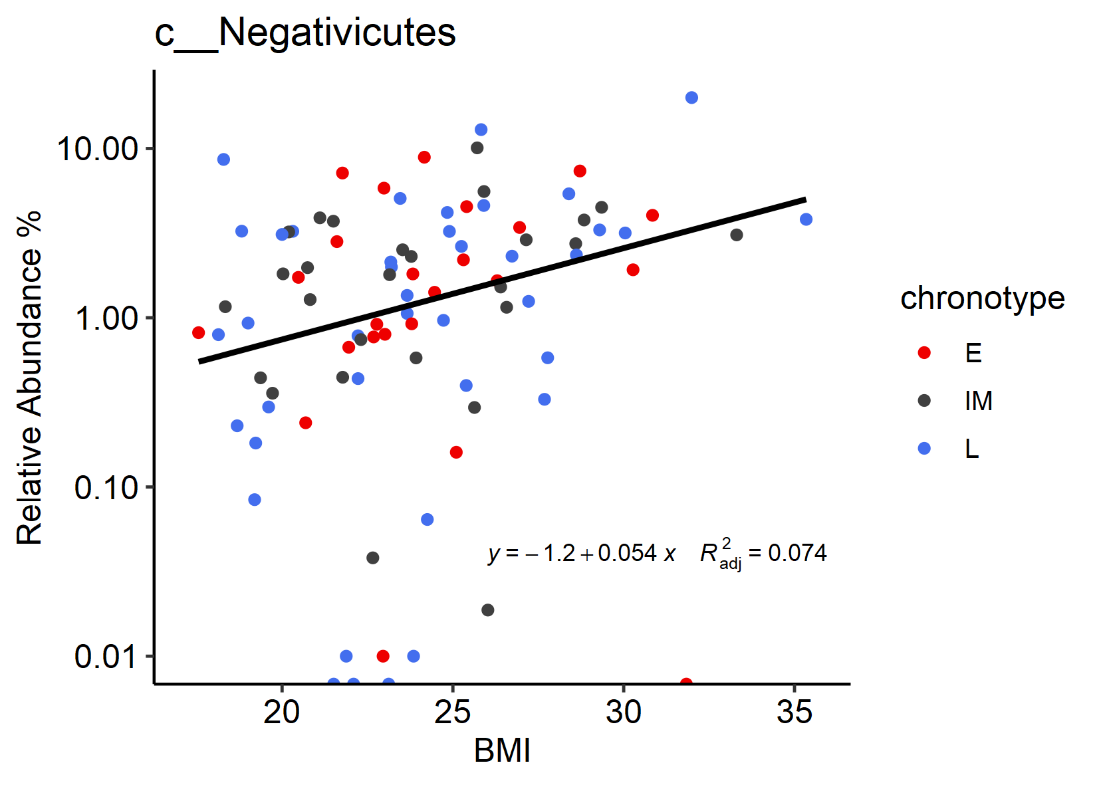
